# Evolution from total variation to nonlinear sparsifying transform for sparse-view CT image reconstruction

**DOI:** 10.1101/785261

**Authors:** Jian Dong, Chunxiao Han, Zhuanping Qin, Yanqiu Che

**Author notes:** Correspondence (Y.C.).

## Abstract

Sparse-view CT has been widely studied as an effective strategy for reducing radiation dose to patients. However, the conventional image reconstruction algorithms, such as filtered back-projection method and classical algebraic reconstruction techniques, can no longer be competent in the image reconstruction task of sparse-view CT. A new principle, called compressed sensing (CS), has been therefore developed to provide an effective solution for the ill-posed inverse problem of sparse-view CT image reconstruction. Total variation (TV) minimization, which is most extensively studied among the existing CS techniques, has been recognized as a powerful tool for dealing with this difficult problem in image reconstruction community. However, in recent years, the drawbacks of TV are being increasingly reported, which are appearance of patchy artifacts, depict of incorrect object boundaries, and loss in image textures or smooth intensity changes. These degradations appear especially in reconstructing actual CT images with complicated intensity changes. In order to address these drawbacks, a series of advanced algorithms using nonlinear sparsifying transform (NLST) have been proposed very recently. The NLST-based CS is based on a different framework from the TV, and it achieves an improvement in image quality. Since it is a relatively newly proposed idea, within the scope of our knowledge, there exist few literatures that discusses comprehensively how the image quality improvement occurs in comparison with the conventional TV method. In this study, we investigated the image quality differences between the conventional TV minimization and the NLST-based CS, as well as image quality differences among different kinds of NLST-based CS algorithms in the sparse-view CT image reconstruction. More specifically, image reconstructions of actual CT images of different body parts were carried out to demonstrate the image quality differences.

## Introduction

Radiation exposure in X-ray Computed Tomography (CT) examinations has raising growing concerns from patients, radiologists, and medical physics communities [1, 2]. Many approaches have been taken to reduce radiation dose of CT scans, including formulation of new hardware-based scanning protocols [3,4] and development of innovative software-based image reconstruction algorithms [5–8]. Sparse-view CT has been widely studied as an effective strategy for reducing radiation dose. Compared with the existing CT systems where several hundred or over a thousand of projection views are required per rotation, as shown in Fig. 1, the sparse-view CT has a potential to reduce the number of projection views to 1/10 of the number employed in current commercial CT scanners. Thanks to the reduction in projection views, patient radiation dose is also significantly decreased. However, due to the insufficient sampling with sparse-view measurements, neither conventional filtered back-projection (FBP) method nor classical algebraic reconstruction technique (ART) method can achieve sufficient diagnostic image quality any more.

**Figure 1:**
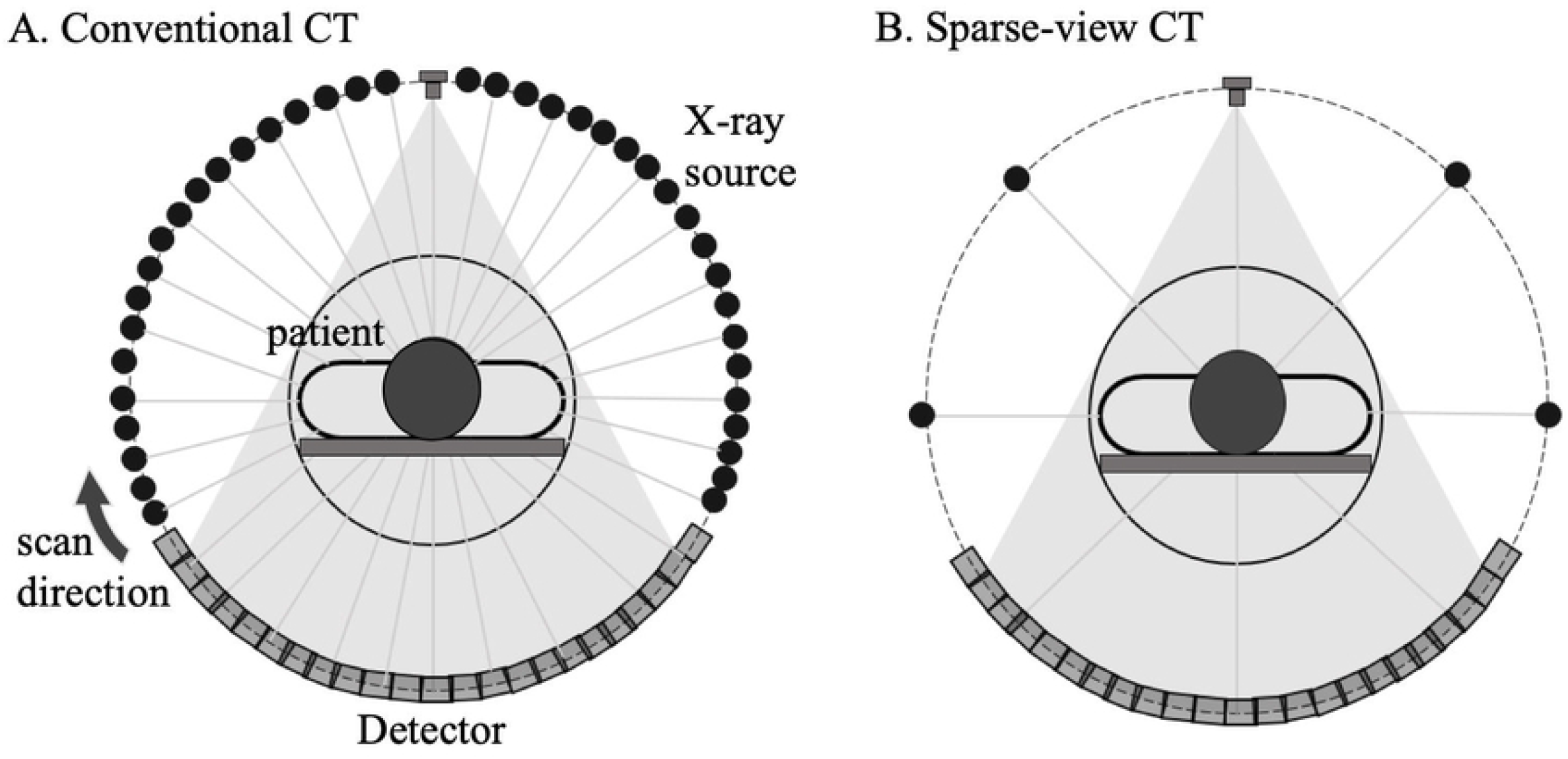
SConfigurations corresponding to two CT imaging situations, i.e. (left) the conventional full-scan CT and (right) sparse-view CT.

Two old works in the mathematical literature demonstrated that the solution to the image reconstruction from a small number of projection data is not unique [9, 10]. Therefore, some prior knowledge on the object needs to be employed to achieve satisfactory reconstructions. Actually, many researchers have investigated this problem with a variety of reconstruction methods and prior knowledge [11, 12], but no one succeeded in discovering an excellent method which can be used in clinical routine. The situation has changed dramatically during 2000’s thanks to the discovery of compressed sensing (CS) theory. The CS was originally proposed by Donoho [13] and Candes et al. [14], and now it has become the gold standard to solve a class of difficult inverse problems including the sparse-view CT image reconstruction. The CS provides an innovative framework to solve the underdetermined ill-posed inverse problems. Generally, a cost function consisting of data fidelity term and penalty term called regularization is designed. Then, an optimal solution is obtained by minimizing the cost function based on convex optimization techniques. The data fidelity term of CS is regarded as the fundamental part, and the least-squares error between estimated projection data and actually measured projection data is commonly used. The regularization plays an important role to compensate for missing information in the incomplete sparse-view projection data, where different regularizations lead to different solutions.

Total variation (TV) is attracting much attention as a kind of regularization used in the CS, because it has achieved big success in producing high image quality in the task of sparse-view CT reconstruction. The TV is defined as a measure to evaluate sum of the intensity gradient at each pixel, and it was first proposed by Rudin, Osher and Fatemi [15] in 1992. In early stages, the TV was used as a penalty term in iterative CT image reconstruction algorithms [16–19], with significant success in smoothing out statistical noise in the image while preserving sharp object boundaries. Subsequently, Sidky et al. extended the application of TV to a constrained optimization approach, where the TV was a cost function and data fidelity was a constraint [20,21]. In this case, they found the solution minimizing the TV under the hard constraint corresponding to the data fidelity. This extension made the TV approach to be more robust in treating the underdetermined problem such as sparse-view and limited angle reconstruction. Lately, a further modification of the TV, i.e. weighted-TV, has been proposed in which they took the edge property of natural images into account [22,23]. This proposal has a potential to avoid the excessive smoothing boundaries occurred in the TV to some extent. The TV is becoming the most standard approach of CS in CT image reconstruction. In recent years, however, the drawbacks of TV, such as depict of incorrect object boundaries, and loss in image textures or smooth intensity changes, are being increasingly reported [24–26], which accelerates comprehensive understanding about the limitations of TV. Additionally, a decade ago, Herman and Davidi claimed that the TV favors no intensity changes in images and has limitations in exactly reconstructing images with complicated structures [27]. Furthermore, they reasonably demonstrated the mechanism of TV limitation. Therefore, a strong demand has been raised towards developing a more powerful CS framework for the sparse-view CT image reconstruction.

Recently, several works have launched a new direction in formulating the regularization term [28–31]. It is known that the key process of designing the regularization term is a concept called “sparsity”. A sparsifying transform to the object image is routinely carried out, which enforces the majority of pixel values to be approximately zeros. The conventional TV achieves “sparsity” by using a linear sparsifying transform (LST), i.e. evaluating intensity gradient of each pixel. However, the new idea focuses on achieving “sparsity” by using a nonlinear sparsifying transform (NLST). A class of nonlinear filters, such as median filter, bilateral filter or nonlocal means (NLM) filter, is combined with the regularization term. Based on the results of these new works, it has been gradually concluded that the NLST has superiority in improving image quality in the task of image reconstruction compared with the conventional TV approach. The mechanism is briefly outlined below. The TV approach performs a same degree of smoothing in the sparsifying transform to the whole image by computing the difference between adjacent pixels. On the contrary, NLST such as Nonlocal Means (NLM) and bilateral filters are selecting the spatially-variant filter to be used in the sparsifying transform by taking image properties, i.e. intensity change around each pixel, into account. A strong smoothing is applied when there exist many similar intensities in the neighboring pixels, and a weak smoothing is applied when similar intensities are few in the neighborhood. Therefore, the NLST can be regarded as a kind of spatially-varying processing, which should contribute to improving image quality.

Based on the aforementioned analysis, superiority of the NLST compared with the TV is gradually becoming clear so that the NLST-based CS is a powerful evolved algorithm. However, to the best of our knowledge, no works have shown a significant image quality difference between the TV and the NLST-based CS. In this paper, we investigated how image quality differs between the conventional TV approach and the NLST-based CS. Furthermore, we investigated how image quality changes when using various kinds of NLSTs. To demonstrate these issues, actual CT images from different body parts were reconstructed from rather small number of projection data.

## Material and methods

### Problem definition

The aim of CT image reconstruction is to recover an object from measured projection data. When iterative methods are used for image reconstruction, the problem can be formulated as solving a linear equation 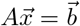, where 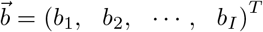 denotes the measured projection data, 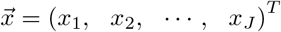 represents the attenuation coefficients of object to be reconstructed, and *A* = {*a*_ij_} is the *I* × *J* system matrix. When the dimension of 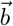 is smaller than the dimension of 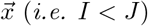, the problem becomes underdetermined so that reconstructing an accurate image becomes challenging, which is the standard mathematical setup of sparse-view CT reconstruction. In such situation, CS is usually utilized to obtain a feasible image, in which a cost function consisting of a data fidelity term 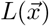 and a regularization term 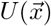 in Eq. (1) is minimized.

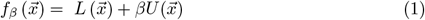

where 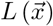 is defined by the least-squares error 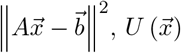 is defined by Eq. (2) below, and *β* is the hyperparameter to control the strength of regularization.

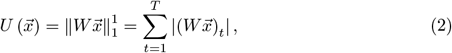

where *W* represents the sparsifying transform which converts 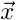 into a sparse vector, and *T* denotes the dimension of sparsifying transformed coefficients vector 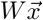. In Eq. (2), the norm 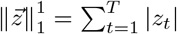 is called *L*1 norm which is known to have superior ability in picking up sparse solutions [6,32]. The sparsifying transform W is usually designed as a high-pass filter which extracts high frequency components of object (*i.e*. object boundaries and textures). In this paper, for the purpose of this study, which is to compare image quality of the TV and the NLST, we design W with two different forms, which correspond to the TV and the NLST, respectively. The detailed formulas are given by Eqs. (3) and (4), where Eq. (3) is the expression of TV case and Eq. (4) is the expression of NLST case.

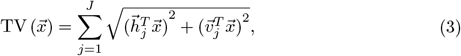

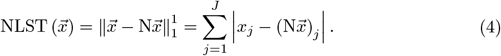

In Eq. (3), 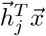 and 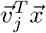 are inner product representations of finite difference operations around the *j*-th pixel along the horizontal and vertical directions, respectively. See Fig. 2 for the detailed definitions of 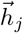 and 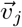. The meaning of Eq. (3) can be interpreted as 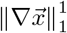, which is the *L*1 norm of the magnitude of intensity gradient for a given image. In Eq. 4, N denotes an arbitrary nonlinear low-pass filter. It is known that nonlinear filters can frequently show satisfactory performance in the task of image processing such as denoising. In this study, we investigate the potential effectiveness of nonlinear filters in the task of image reconstruction, to compare the result of the NLST with that of the TV approach. We utilized median filter, bilateral filter and nonlocal means (NLM) filter as the non-linear filter N to perform the sparsifying transform.

**Figure 2:**
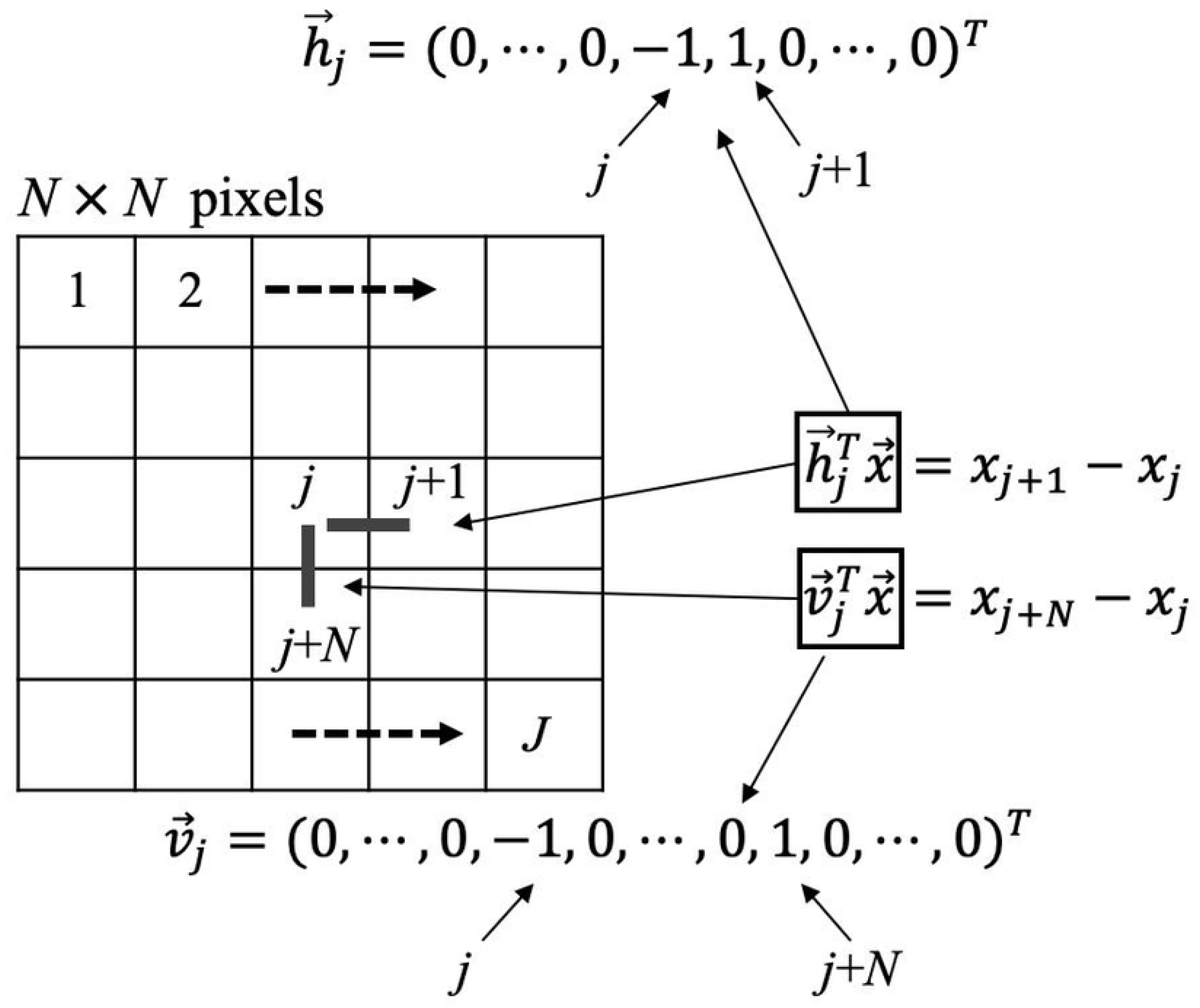
Definitions of the horizontal difference 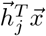 and the vertical difference 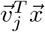 used in the TV.

### Mathematical tools

Once the cost function is determined, the crucial subsequent step is to derive an iterative algorithm to find the optimal solution by minimizing the cost function. We utilize techniques of convex optimization to achieve this task. Before going into the concrete derivation, it is indispensable to introduce two mathematical tools together with some related basic pieces of prior knowledge. Let us consider a convex minimization problem formulated as

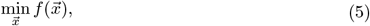

where we assume that 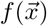 is a possibly non-differentiable lower semi-continuous (lsc) convex function.

[**Proximity Operator and Proximal Algorithm**] [33] The proximity (*prox*) operator corresponding to 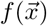 is defined by

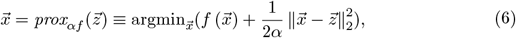

where the parameter *α* is called the step-size parameter. There exist two convenient properties in the *prox* operator, which facilitated us to use this operator in the design of iterative algorithm. One property is that it can be defined for any lsc convex function even if it is non-differentiable like the *L*1 norm. The other property is that its fixed points, *i.e*. 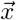 satisfying 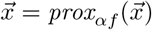, coincide with minimizers of 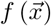 for any *α* > 0. Furthermore, the *prox* operator is necessarily a non-expansive mapping. These properties allow us to use the following iteration formula to find a minimizer of 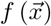.

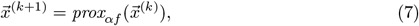

where *k* denotes the iteration number. This iterative algorithm is called the proximal minimization algorithm, which provides a powerful framework for non-differentiable convex minimizations.

[**Proximal Gradient Method**] Let us consider a convex minimization problem expressed as

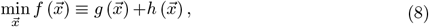

where 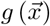 is a smooth convex function and 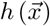 is a possibly non-differentiable convex function. We consider the situation where the *prox* operator 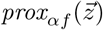 corresponding to 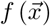 is intensive to compute, but 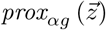 and 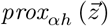 corresponding to the sub-cost functions 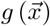 and 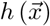 can be easily computed. The proximal gradient method is a framework for constructing an iterative algorithm to minimize 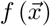 under such situations [34]. Basically, the algorithm is constructed by first computing an intermediate variable 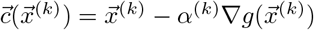 by implementing gradient descent processing to the smooth function 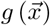. And then achieve the updated vector 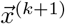 by computing the *prox* operator 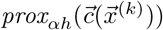 corresponding to 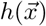. The processing procedure can be summarized as

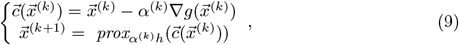

where *k* denotes the iteration number, and *α*^(*k*)^ is the step-size parameter dependent on *k*. With respect to the convergence property of Eq. (9), Passty proved the following theorem [35], where a more general situation of multiple, *i.e*. more than two, sub-cost functions were considered. Then the following theorem holds.

**Theorem** We consider an ergodic average of the iterates 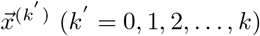 defined by

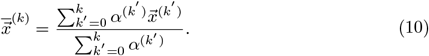

Then, 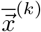 converges to a minimizer of 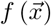 when the diminishing step-size who satisfies the following rules is used.

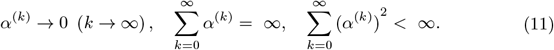

We note that the above convergence is called the ergodic convergence, in which the ergodic average converges to a minimizer of 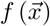 more easily than the sequence 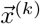 itself. In practice, however, we have never observed a situation in which the ergodic convergence occurs with the sequence 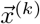 itself being not convergent. Therefore, we conjecture that the proposed algorithm expressed by Eq. (9) can be practically used without being anxious about the non-convergence.

### Iterative algorithm for total variation reconstruction

In this section, we explain how to construct the iterative algorithm for the TV minimization by using the proximal gradient framework. The cost function of TV can be expressed in detail by

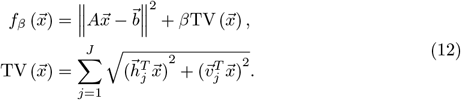

We first split the cost function in Eq. (12) into the sum of sub-cost functions as

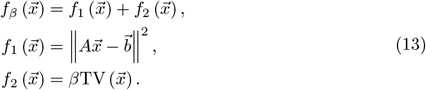

Then the gradient descent processing and *prox* operator can be applied to each sub-cost function alternately, which leads to the two-step iterative algorithm as

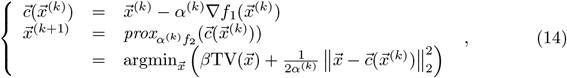

where *k* denotes the iterative number. The algorithm expressed by Eq. (14) still does not have a concrete implementable form. Thus, we make further simplification. To achieve the intermediate image 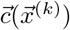 from the first equation in Eq. (14), a differential operation to the smooth function 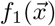 is usually carried out. It is clear that 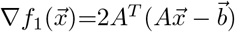. We can obtain the iterative formula expressed as

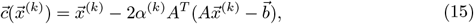

where *A^T^* indicates the transpose matrix of system matrix *A*. The iteration formula in Eq. (15) possesses a well-known structure which is the same as that in the steepest descent method applied to the minimization of 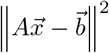.

Next, we deal with the computation of *prox* operator corresponding to the TV term (*i.e*. solving the second equation of Eq. (14)). This minimization problem is the same as that appearing in the so-called TV denoising (ROF model), so that we can use a standard algorithm such as Chambolle’s projection algorithm [15,36]. In our implementation of image reconstruction, we used Chambolle’s projection algorithm. We give a detail explanation about this algorithm in Appendix A.

In order to guarantee the convergence of the proposed algorithm, we take a commonly used way to set the step-size parameter *α*^(*k*)^ as

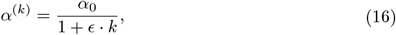

where *α*_0_ > 0 and *ϵ* > 0 are pre-specified parameters, and *k* denotes the iteration number.

So far, the iterative algorithm for the TV reconstruction can be summarized in Algorithm 1.

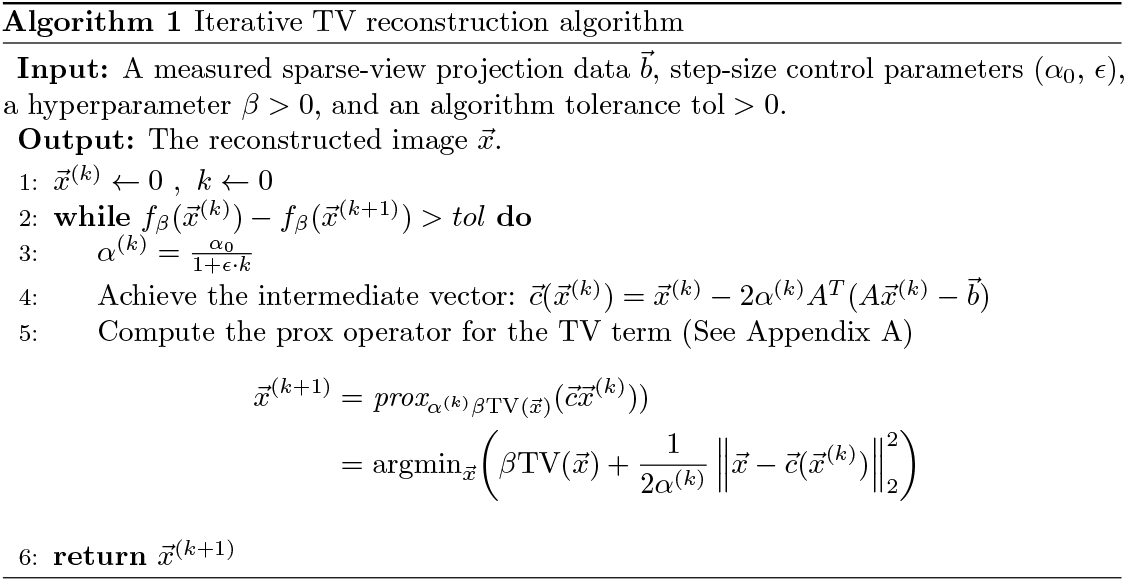

### Iterative algorithm for nonlinear sparsifying transform reconstruction

In this section, we explain how to construct the iterative algorithm for the NLST by using the proximal gradient framework. Based on the derivation in Section 2.1, the cost function of NLST can be expressed as

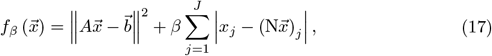

where N denotes a kind of nonlinear low-pass filter. In this study, we utilize three typical nonlinear filters, which are median, bilateral and NLM filters. The relatively simple median filter first sorts pixel intensities inside neighboring window and replaces the central pixel intensity by the median value of the pixel intensities. Both the bilateral filter and the NLM filter replace the central pixel intensity by the weighted average of pixel intensities inside the neighboring search window. The bilateral filter computes the weights by evaluating distance and intensity similarities between a single pixel located within search window and the central pixel, while NLM filter computes the weights based on the evaluation of specific patch widows. Eq. (18) shows the common computation formula corresponding to the two filters. Eqs. (19) and (20) show the specific weight computation method for the bilateral and the NLM filter, respectively.

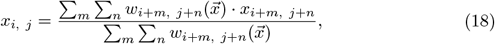

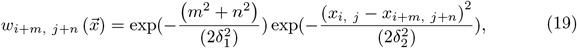

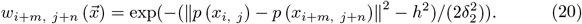

Note that, to simplify the expressions in Eqs. (18)–(20), we used two-dimensional representation style for each pixel, where (*i, j*) denotes the coordinates of the central pixel, and (*m, n*) denotes the coordinates of the search window. The *δ*_1_ controls the weight of Gaussian distance, *δ*_2_ controls the weight of intensity similarity, *h* denotes the constant to take the statistical noise into account, and ||p(*x_i, j_*) − p(*x*_*i*+*m*, *j*+*n*_)||^2^ is the Euclidean intensity distance between two different patches which are centered at pixel *x_i, j_* and *x*_*i*+*m*, *j*+*n*_, respectively.

Similarly to the derivation in Section 2.3, we split the cost function into the sum of two sub-cost functions as

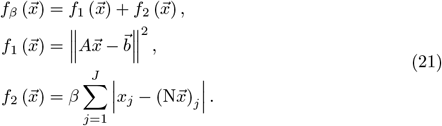

Then, the gradient descent processing and *prox* operator can be applied to the sub-cost functions alternately, and we can obtain the two-step iterative algorithm as

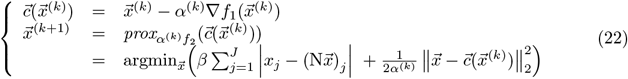

where *k* denotes the iteration number. We observe that the first equation in Eq. (22) is exactly the same as that in Eq. (14), thus we can use the same iteration formula as in Eq. (15). Next, we explain the solution of the *prox* operator corresponding to the NLST term (the second equation in Eq. (22)). It is a function consisting of an absolute value term and a quadratic term. The minimization of this function can be approximately performed in a closed form by using the so-called the soft-thresholding operation as follows. First, we note that there exists an obstacle to apply the soft-thresholding operation in this minimization, because the term 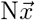 is dependent on the variable 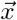. Here, we use a trick called “constant approximation”. When the step-size *α*^(*k*)^ is set to a sufficiently small value, the unknown 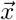 approximates infinitely close to the intermediate image 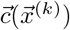, and it is expected that 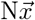 is approximated well by 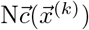, which succeeds in converting the absolute value term into a separable form. Thanks to this approximation, an iterative formula expressed by Eq. (23) can be obtained by applying the soft-thresholding operation.

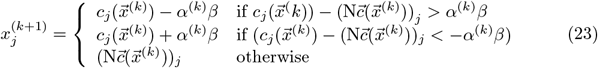

With respect to the control of step-size parameter *α*^(*k*)^, we use the same method as in the TV approach, *i.e*. *α*^(*k*)^ is decreased toward zero according to the diminishing step-size rule in Eq. (16).

In addition, we make the following modification into the algorithm to further improve the performance. For the NLST algorithm, some literatures have shown that, when it is used in image reconstruction or image recovery [28, 37], the NLSM can cause isolated points in the image. The reason is explained as follows. The nonlinear filters, such as the NLM and bilateral filters, determine the spatially-dependent degree of smoothing based on the image characteristic at the current iteration. A strong smoothing is applied when there exist many similar intensities in the neighboring pixels. On the contrary, a weak smoothing is applied when similar intensities are few in the neighboring pixels. Thanks to the spatially dependent nature of sparsifying transform, the NLST strengthens its ability to improve image quality compared with the TV approach. However, the filter is determined by using the intermediate images containing heavy streak artifacts during iterations. The choice of inaccurate smoothing filter or fairly weak smoothing filter often causes to generate isolated points in the image. On the other hand, the TV approach performs the same degree of smoothing by computing the differences between adjacent pixels at every pixel location. As this procedure is not spatially-dependent, the TV does not cause the isolated points. Therefore, in order to prevent the NLST algorithm from suffering from the isolated points, we add the TV regularization with a tiny weight working only on eliminating the isolated points to the NLST regularization term.

Finally, we can summarize the iterative algorithm of NLST approach as in Algorithm 2.

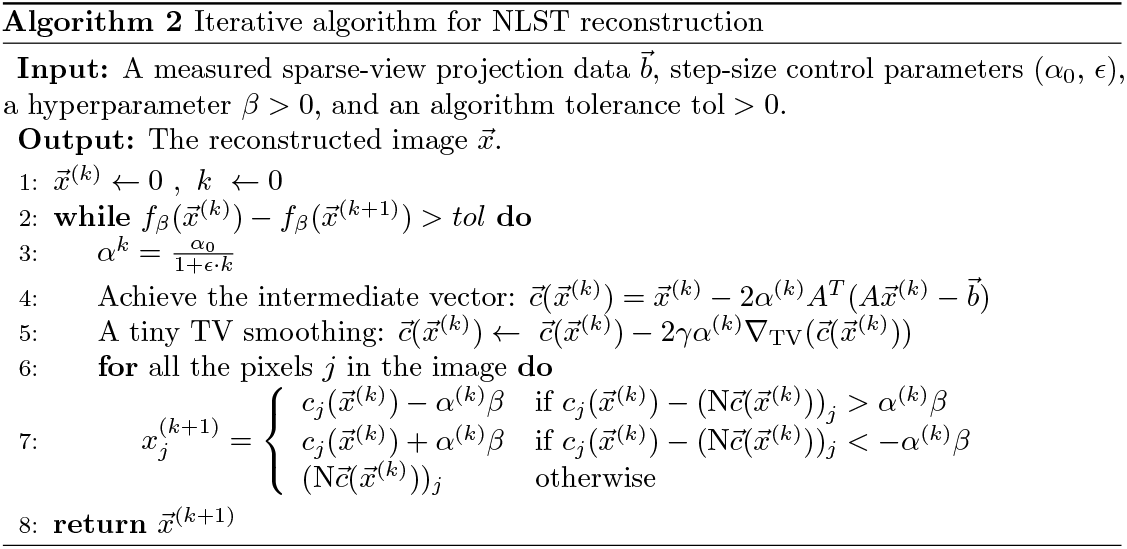

## Experimental results

In this section, we investigated effectiveness of the conventional TV method and the NLST method for the task of image reconstruction in sparse-view CT. In this study, we carried out image acquisition from medical image library of *The Japanese Society of Medical Imaging Technology*. The images in this library are provided for medical image processing. The image data format was DICOM (Digital Imaging and Communications in Medicine), but the patient content was removed in advance. Thus our study did not involve human participants. Before the study began, we obtained the approval from ethics committee of *School of Automation and Electrical Engineering, Tianjin University of Technology and Education*. Since CT examinations of upper abdominal part are routinely carried out in clinical diagnosis, we first applied the methods to the reconstruction of a CT trans-axial slice containing liver and spleen. The original CT image used in the simulation consists of 512 × 512 pixels, *i.e*. pixel size is 320 (mm) × 320 (mm). As a comparison, in Fig. 3, we show a reconstructed image from 1000 projection views (over the angular range of 180 degrees) by the conventional FBP method. In Fig. 4, we show reconstructed images from only 72 projection views (over the angular range of 180 degrees) by the TV and NLST methods. Fig. 4(a) is the reconstructed image with 1000 iterations by the TV method. The used implementation parameters are summarized as follows: *α*_0_=100, *ϵ*=100 and the hyperparameter *β* = 50. Fig. 4(b-d) show reconstructed images by the NLST method. In this method, we set the hyperparameter as *β* = 35, and the step-size control parameters were *α*_0_=100 and *ϵ*=1000. Fig. 4(b) is the reconstructed image with 1000 iterations by the NLST method using the median filter. We set the window size of median filter to 3× 3 pixels. Fig. 4(c) is the reconstructed image with 1000 iterations by the NLST method using the bilateral filter. The implementation parameters to obtain this image were summarized as follows: window size is 11×11 pixels, *δ*_1_=10000, and *δ*_2_=120. Fig. 4(d) is the reconstructed image with 1000 iterations by the NLST method using the NLM filter. The parameters of the NLM filter were set as follows: search window size is 7× 7 pixels, patch window size is 5×5 pixels, *δ*_1_=15, and *δ*_2_=15. In Fig. 4, the small parts of the images were zoomed-in and displayed on the left or right side. Significant image degradations can be obviously confirmed on the reconstructed image of Fig. 4(a) by the TV method. In particular, severe patchy artifacts can be clearly observed, and the hepatic veins were partly lost. Experimental results of Figs. 4(b-d) demonstrate that the NLST method can undoubtedly improve image quality with respect to reducing the patchy artifacts, preserving object boundaries accurately and image textures. Also, we can observe that using different nonlinear filters in the regularization term of the NLST method can lead to different image quality. The NLST method using the median filter produced the images with still remaining patchy artifacts, the NLST method using the bilateral filter depicted image textures with somewhat serrated boundaries, and the NLST method using the NLM filter achieved the best image quality.

**Figure 3:**
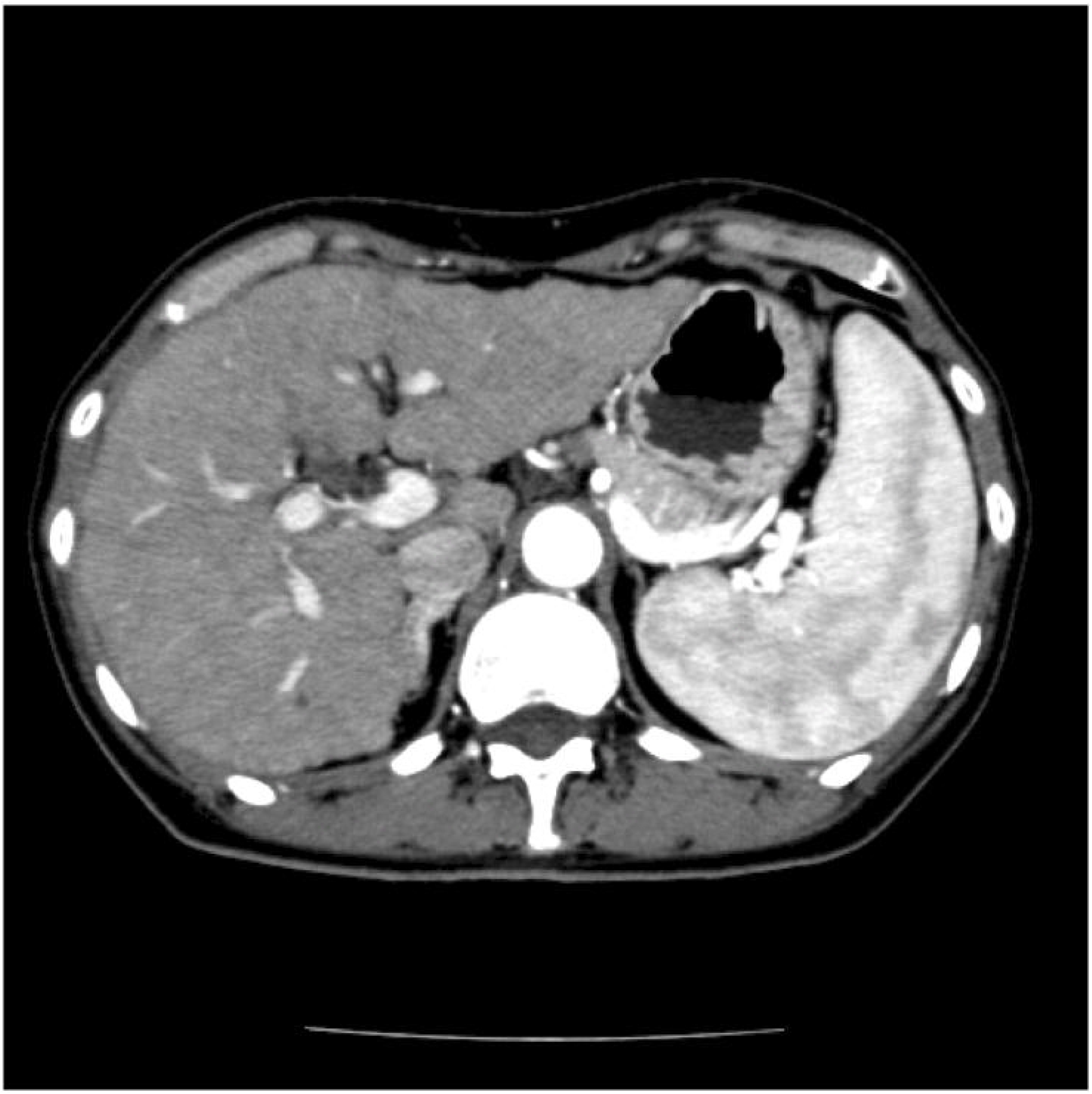
Reconstructed image by the standard FBP method with 1000 projection views (over the angular range of 180 degrees).

**Figure 4:**
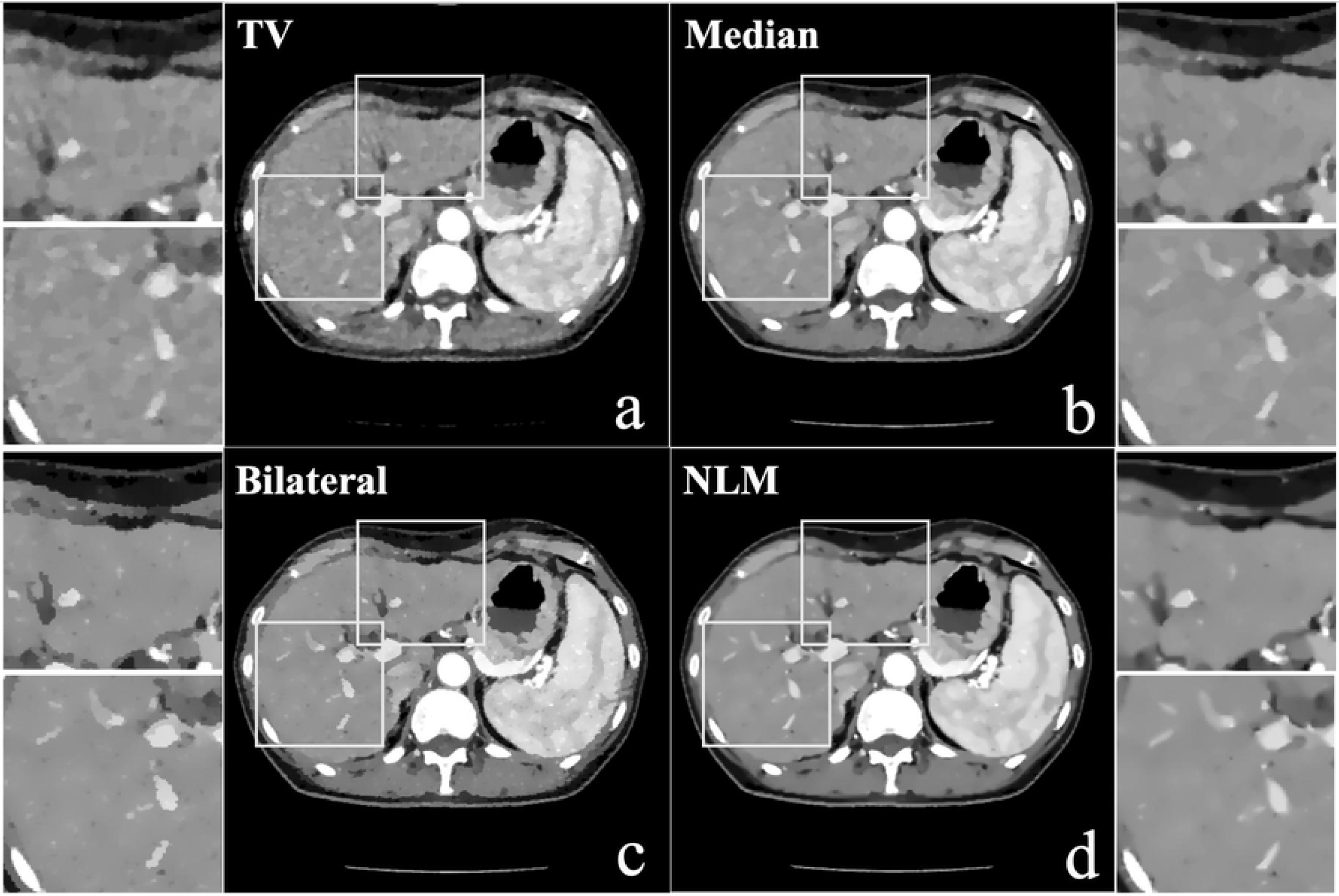
Reconstructed images by the conventional TV method (a) and the NLST methods (b: median filter, c: bilateral filter, d: NLM filter) from 72 projection views (over the 180 angular range of 180 degrees).

In Fig. 5, we show reconstructed images by the conventional TV method with different hyperparameter values *β*. Since *β* controls the strength of regularization, the different values of *β* can lead to images with different degree of smoothing. Thus, we investigated effectiveness of the TV method with different *β* values. Similarly to the experiments in Fig. 4, 72 projection views were used for image reconstruction, and the step-size control parameters were set to *α*_0_=100 and *ϵ*=100. Also, 1000 iterations were carried out in this experiment. When a small hyperparameter value of *β*=1.0 was used, the strength of TV was so light that the streak artifacts (the part pointed by the arrow) could not be completely removed. On the contrary, when the too large hyperparameter value of *β*=100.0 was used, the reconstructed image was forced to be close to be piecewise constant, which excessively smoothed the image textures (the part pointed by the arrow) and some of the textures were lost. As observed from this result, the TV method has limitations when applied to this severely ill-posed inverse problem.

**Figure 5:**
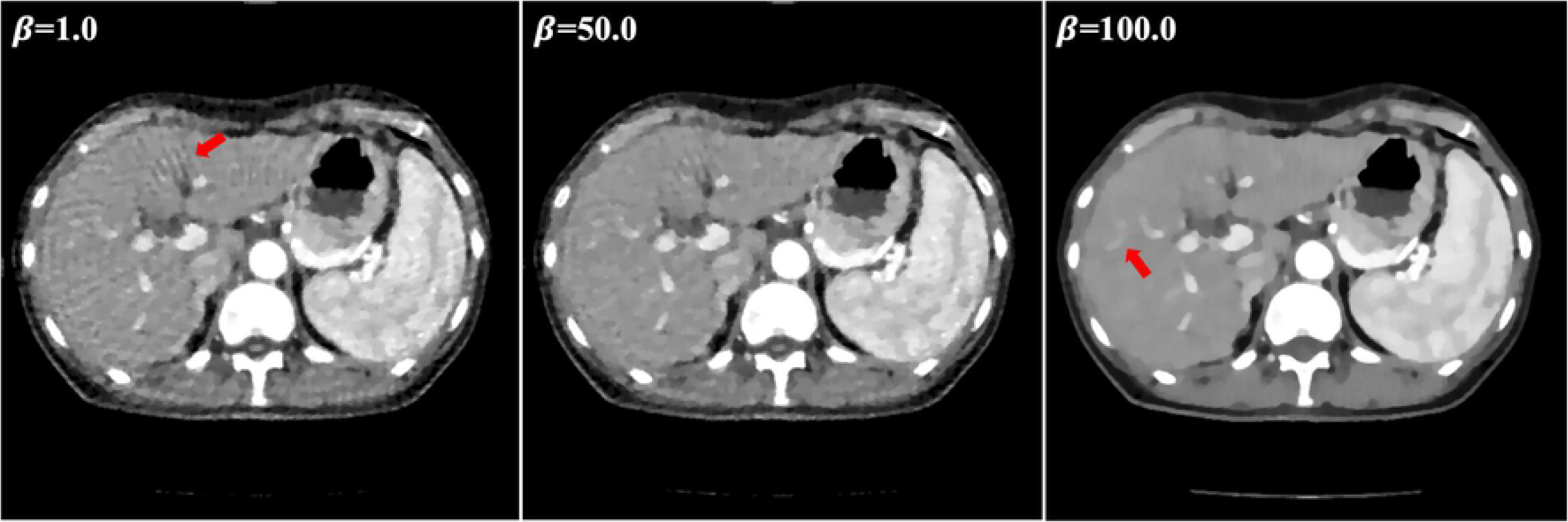
Reconstructed images by the TV method with three different values of hyperparameter *β*.

Considering that image characteristics of CT images differ significantly dependent on parts of human body, we applied the described methods to reconstruct CT images for four other body parts, which were dental area, cardiac area, liver area, and abdominal area. The pixel size of each original image was 512 × 512. We displayed the experimental results in Figs. 6–9. More specifically, Fig. 6 shows reconstruction results of dental part from only 32 projection views, Fig. 7 shows reconstruction results of cardiac part from only 32 projection views, Fig. 8 shows reconstruction results of liver part from only 32 projection views, and Fig. 9 shows reconstruction results of abdominal part from only 48 projection views. For comparison, on the left of each figure, we also provided an image reconstructed by the standard FBP method with 1000 projection views. Then, we compared image quality difference between the TV method and the NLST method. In the implementation of NLST method, we employed the NLM filter, because the above-mentioned experimental result demonstrated that it is the most successful nonlinear filter in producing a high-quality image. With respect to the parameter settings in implementing the TV method and the NLST method, we show a detailed summary of the used parameter values in Table 1. The reconstruction results for the four body parts indicated the same tendency as the first study of abdominal CT image case, *i.e*. the NLST method could improve image quality significantly compared with the TV method. The reconstruction results of each body part by the TV method suffered from heavy patchy artifacts, but they were successfully reduced by the NLST method. Besides, the reconstructed images by the NLST method preserved image textures and object boundaries accurately.

**Figure 6:**
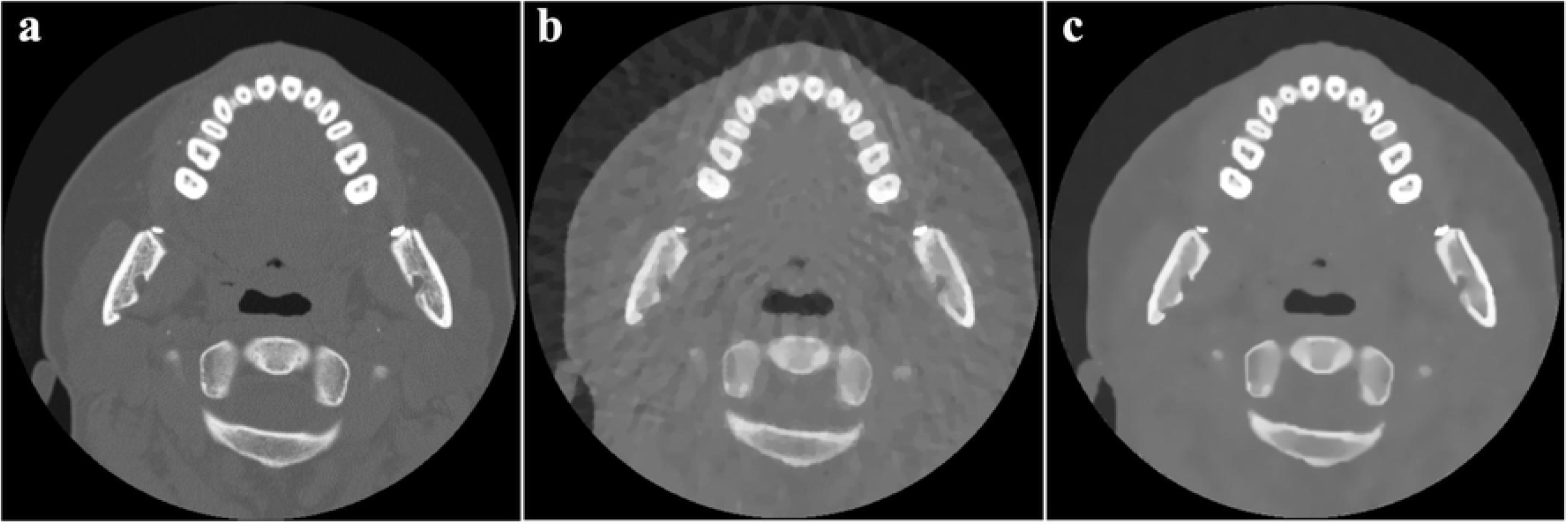
Reconstructed images for the dental CT image by (a) the FBP method with 1000 projection views, (b) the conventional TV method with 32 projection views, and (c) the NLST method using the NLM filter with 32 projection views.

**Figure 7:**
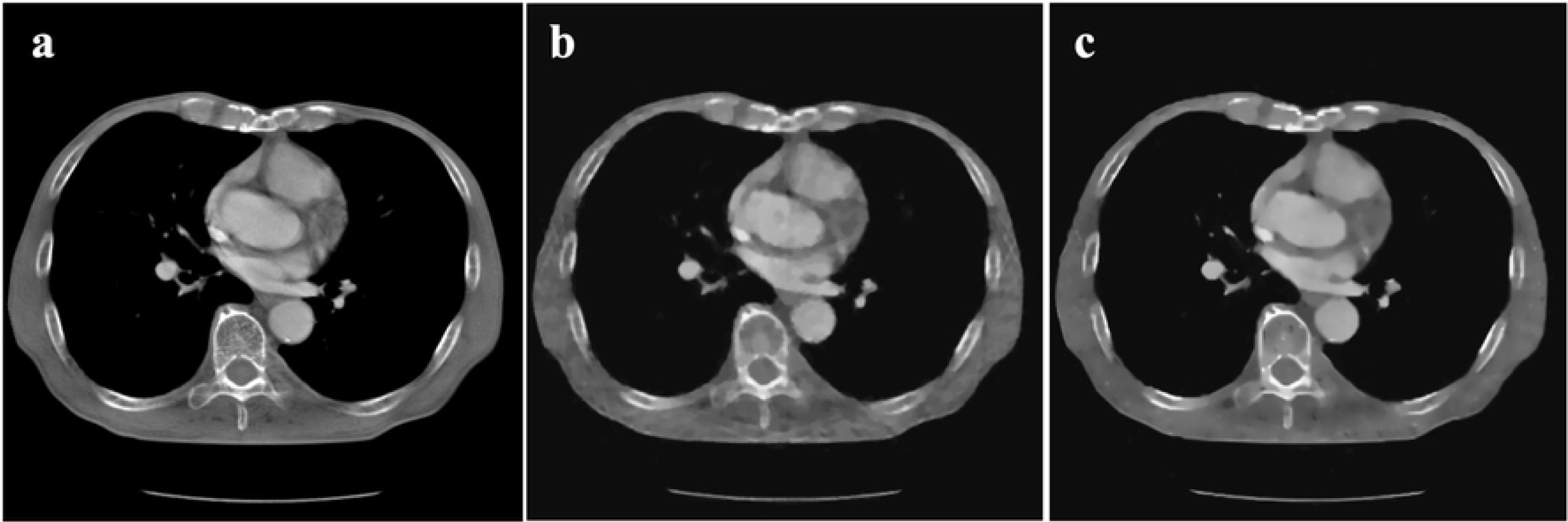
Reconstructed images for the cardiac CT image (a) by the FBP method with 1000 projection views, (b) the conventional TV method with 32 projection views, and (c) the NLST method using the NLM filter with 32 projection views.

**Figure 8:**
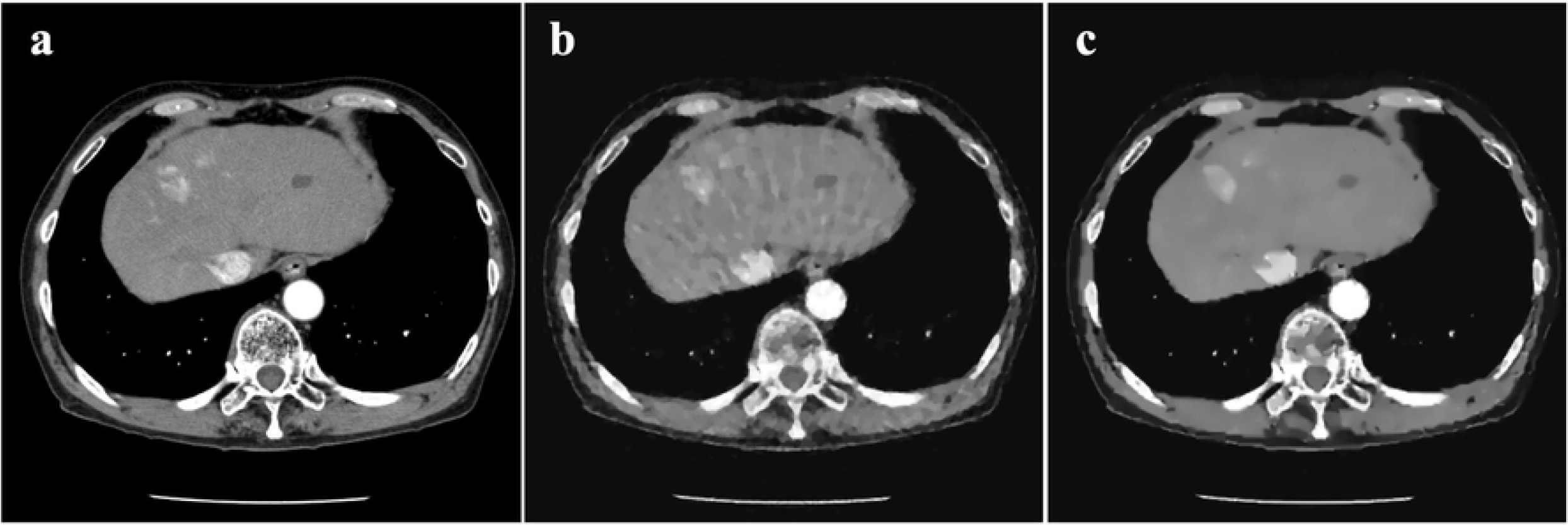
Reconstructed images for the hepatic CT image (a) by the FBP method with 1000 projection views, (b) the conventional TV method with 32 projection views, and (c) the NLST method using the NLM filter with 32 projection views.

**Figure 9:**
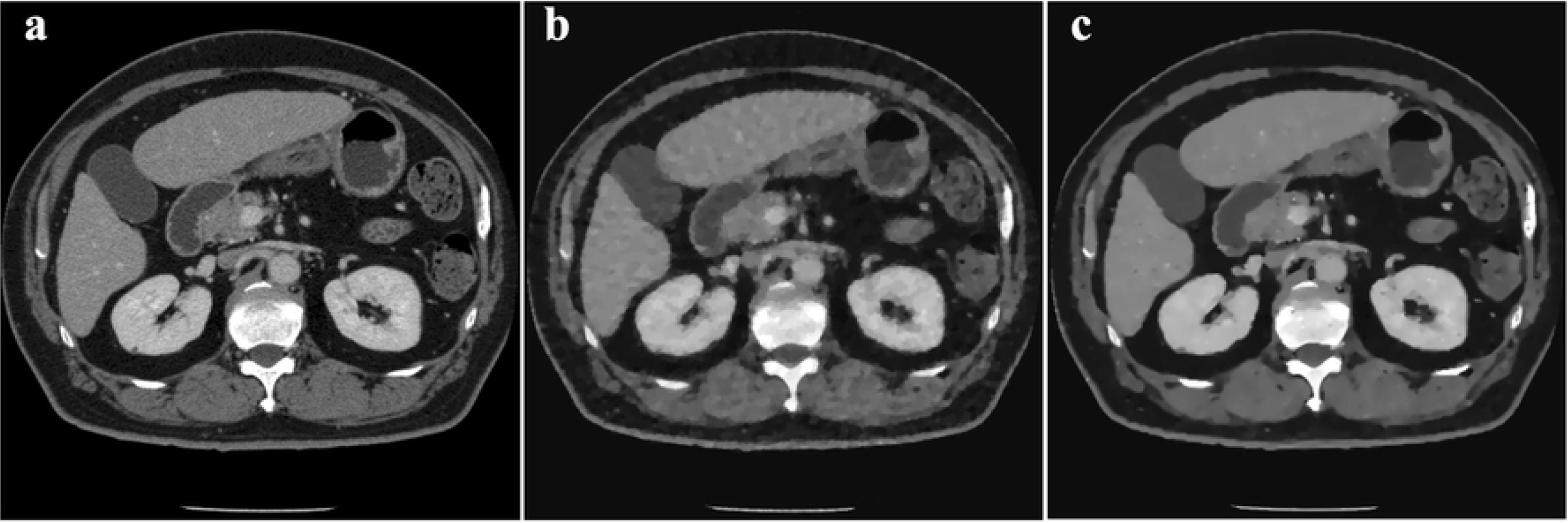
Reconstructed images for the abdominal CT image (a) by the FBP method with 1000 projection views, (b) the conventional TV method with 48 projection views, and (c) the NLST method using the NLM filter with 48 projection views.

**Table 1.** Parameter values in respective reconstruction of practical CT image instances in TV and NLST methods.

|  |  | Dental CT | Cardiac CT | Hepatic CT | Abdominal CT |
| --- | --- | --- | --- | --- | --- |
| TV | $\alpha_0$ | 100 | 100 | 100 | 100 |
| | $\epsilon$ | 100 | 100 | 100 | 100 |
| | $\beta$ | 65 | 100 | 100 | 100 |
| NLST | $\alpha_0$ | 100 | 100 | 100 | 100 |
| | $\epsilon$ | 1000 | 1000 | 1000 | 1000 |
| | $\beta$ | 70 | 100 | 170 | 80 |
| | $\gamma$ | 1.3 | 1.3 | 5.0 | 4.3 |
| | $\delta_1$ | 5.0 | 25 | 20 | 20 |
| | $\delta_2$ | 5.0 | 25 | 10 | 20 |
| | Window Size | $5 \times 5$ | $5 \times 5$ | $5 \times 5$ | $5 \times 5$ |

## Discussion and conclusion

In this paper, we implemented the conventional TV minimization algorithm and the NLST-based CS algorithm. The implementations were carried out for image reconstruction in sparse-view CT. The TV minimization was achieved by the Chambolle’s projection algorithm. The nonlinear filters including median filter, bilateral filter and NLM filter were respectively embedded into the NLST-based CS algorithm. Actual trans-axial CT slices from different body parts, such as dental area, cardiac area, abdominal area, and hepatic region, were reconstructed for comparison between the two methods. We investigated the resulting image quality differences between the TV and NLST methods, as well as image quality differences induced by different nonlinear filters used in the NLST method.

As observed from the experimental results, the TV method produced degraded images under the same sparse-view CT measurement conditions. When the hyperparameter *β*, which controls the strength of TV term, is too small, the image degradations induced by the TV method include patchy artifacts, streak artifacts, and inaccurate object boundaries. On the contrary, when *β* is too large, the image degradations occur by producing regions having piecewise constant intensity changes and losing image textures and smooth intensity changes. On the other hand, however, the TV method sometimes plays a crucial role in removing isolated points in the reconstructed images, and we used this favorable property of the TV in Algorithm II of this paper. It became clear that the NLST-based CS method can overcome the image degradations of the TV method well. It can produce images which are closer to images obtained by the FBP method with sufficient projection views. Additionally, the NLST-based CS method possesses a potential to embed an arbitrary nonlinear filter into the algorithm. It means that we can extend its application to a specific imaging task. For example, if we are successful in developing a new nonlinear filter which can selectively remove streak artifacts, the NLST-based CS method would contribute to further reducing the number of projection views. In summary, we conclude that the NLST-based CS method is superior to the TV method in the task of image reconstruction for sparse-view CT.

## Appendix A

Here, we give a detailed explanation on how to solve the minimization problem expressed in Eq. (A.1). This minimization problem is the same as the so-called TV denoising (ROF model), and the classical algorithm to solve this problem is Chambolle’s projection algorithm which we describe below.

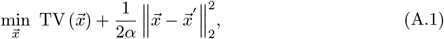

where the vector 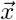 denotes an unknown image with *N* × *N*, and the vector 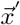 denotes an input image with *N* × *N* contaminated with noise. We use the symbol ∇ to represent the intensity gradient operator, which is defined by

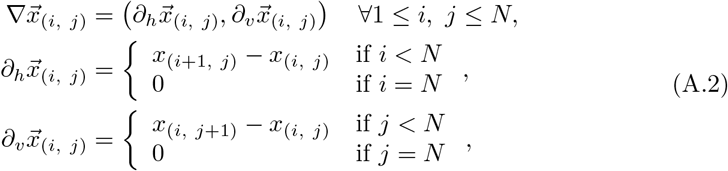

where 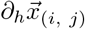 and 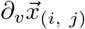 denote the intensity gradient of horizontal and vertical direction, respectively. We introduce two dual variables 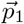 and 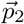, both of which are image vectors with *N* × *N* pixels. Then, we define the divergence operator div by

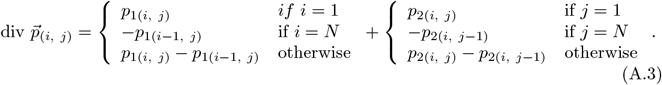

Then, the solution to (A.1) can be obtained by Chambolle’s projection algorithm summarized in Algorithm 3 below.

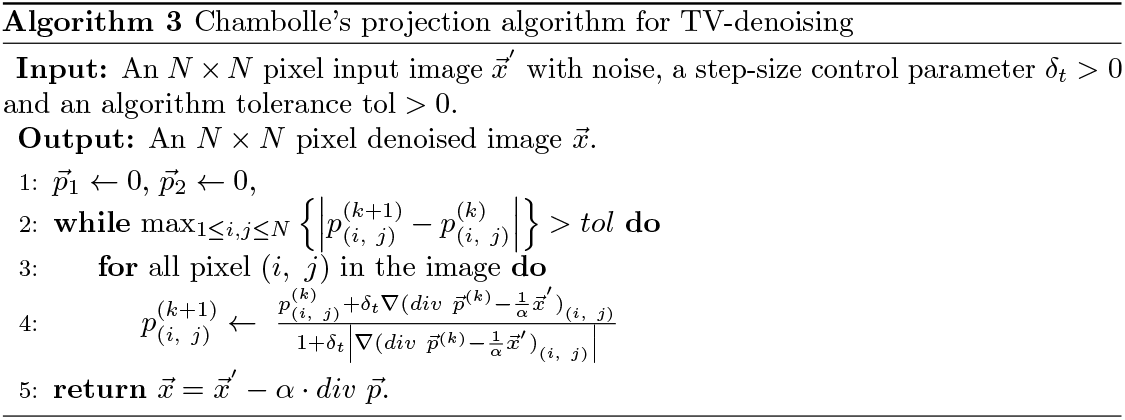

With respect to the convergence of the above algorithm, it was proved that the convergence to the optimum solution is guaranteed when the step-size control parameter *δ_t_* is set to a value smaller than 1/4 [24].

## Acknowledgments

We express our sincere thanks to Prof. Hiroyuki Kudo for his patient teaching on constructing the theoretical framework, and thanks to Ms. Humii Takahama for her assistance in text proofreading. This work was partially supported by Natural Science Foundation of Tianjin City project (Grant No. ZZSK021810) and Tianjin Overseas Returnees Research Funding Support Project (Grant No. 201819) and Tianjin Municipal Special Program of Talents Development for Excellent Youth Scholars (Grant No. TJTZJH-QNBJRC-2-21).

## Author contributions statement

J.D. and Y.C. conceived the research and experiments; J.D., C.H. and Z.Q. conducted the experiments, J.D., C.H., Z.Q. and Y.C. analyzed the results. J.D. and Y.C. wrote the manuscript with critical review by other authors.

## References

1. Brenner DJ, Hall EJ. CT-an increasing source of radiation exposure. New England J Med. 2007;357:2277–2284.

2. Cohnen M, Wittsack HJ, Assadi S, Muskalla K, Ringelstein A. Radiation exposure of patients in comprehensive computed tomography of the head in acute stroke. AJNR Am J Neuroradiol. 2006;27:1741–1745.

3. McCollough CH, Primark AN, Braun N, Kofler J, Yu L, et al. Strategies for reducing radiation dose in ct. Radiol Clin North Am. 2009;47:27–40.

4. Smith A, Dillon W, Gould R. Radiation dose-reduction strategies for neuroradiology ct protocols. Am J Neuroradiol. 2007;28:1628–1632.

5. Kudo H, Nemoto T. Image reconstruction method in interior tomography. In: Patent Application in Japan;.

6. Li M, Yang H, Kudo H. An accurate iterative reconstruction algorithm for sparse objects: application to 3D blood vessel reconstruction from a limited number of projections. Phys Med Biol. 2002;47:2599–2609.

7. Rampinelli C, Origgi D, Bellomi M. Low-dose CT: technique, reading methods and image interpretation. Cancer Imaging. 2013;12:548–556.

8. Wang G, Yu H. The meaning of interior tomography. Phys Med Biol. 2013; p. R161–R186.

9. Louis AK. Ghosts in tomography-The null space of the Radon transform. Math Methods Appl Sci. 1981;3:1–10.

10. Smith KT, Solmon DC, Wagner SL. Practical and mathematical aspects of the problem of reconstructing objects from radiographs. Bull Amer Math Soc. 1977;83:1227–1270.

11. Llacer J. Theory of imaging with a very limited number of projections. IEEE Trans Nucl Sci NS. 1979;26(596-602).

12. Siltanen S, Kolehmainen V, Järvenpää S, et al. Statistical inversion for medical x-ray tomography with few radiographs: I. General theory. Phys Med Biol. 2003;48:1437–1463.

13. Donoho DL. Compressed sensing. IEEE Trans Inf Theory. 2006;52:1289–1306.

14. Candès EJ, Romberg J, Tao T. Robust uncertainty principles: Exact signal reconstruction from highly incomplete frequency information. IEEE Trans Inf Theory. 2006;52:489–509.

15. Rudin LI, Osher S, Fatemi E. Nonlinear total variation noise removal algorithm. Physica D. 1992;60:259–268.

16. Panin VY, Zeng GL, Gullberg GT. Total Variation Regulated EM Algorithm. IEEE Trans Nuc Sci. 1999;46:2202–2210.

17. Persson M, Bone D, Elmqvist H. Total variation norm for three-dimensional iterative reconstruction in limited view angle tomography. Phys Med Biol. 2001;46:853–866.

18. Song J, Liu QH, Johnson GA, Badea CT. Sparseness prior based iterative image reconstruction for retrospectively gated cardiac micro-CT. Med Phys. 2007;34:4476–4483.

19. Vogel CR, Oman ME. Iterative methods for total variation denoising. SIAM J Sci Comp. 1996;17:227–238.

20. Sidky E, Kao C, Pan X. Accurate image reconstruction from few-views and limited-angle data in divergent beam ct. J X-ray Sci Technol. 2006;14:119–139.

21. Sidky E, Pan X. Image reconstruction in circular cone-beam ct by constrained, total-variation minimization. Phys Med Biol. 2008;53:4777–4807.

22. Liu Y, Ma J, Fan Y, Liang Z. Adaptive-weighted total variation minimization for sparse data toward low-dose x-ray computed tomography image reconstruction. Phys Med Biol. 2012; p. 7923–7956.

23. Tian Z, Jia X, Yuan K, Pan T, Jiang SB. Low-dose ct reconstruction via edge-preserving total variation regularization. Phys Med Biol. 2011;56:5949–5967.

24. Cai J, Dong B, Osher S, Shen Z. Image Restoration: Total variation, wavelet frames, and beyond. J Amer Math Soc. 2012;25:1033–1089.

25. Guo W, Qin J, Yin W. A new detail-preserving regularization scheme. SIAM Journal on Imaging Sciences. 2014;.

26. Kudo H, Dong J, Kamo K, Horii N, Furukawa H, et al. Tomographic image reconstruction using compressed sensing. Kenbikyo. 2016;51:48–53.

27. Herman GT, Davidi R. On image reconstruction from a small number of projections. Inverse Probl. 2008;.

28. Dong J, Kudo H. Proposal of Compressed sensing using nonlinear sparsifying transform for CT image reconstruction. Med Imag Tech. 2016;34:235–244.

29. Li Z, Yu L, Trzasko J, Lake D, Blezek D. Adaptive nonlocal means filtering based on local noise level for CT denoising. Med Phys. 2014;41:011908.

30. Lou Y, Zhang X, Osher S, Bertozzi A. Image recovery via nonlocal operators. J of Scientific Computing. 2010;42:185–197.

31. Zhang H, Ma J, Wang J, et al. Statistical image reconstruction for low-dose CT using nonlocal means-based regularization. Comput Med Imaging Graph. 2014;38:423–435.

32. Elad M. Sparse and Redundant Representations: From Theory to Applications in Signal and Image Processing. Springer; 2010.

33. Parikh N, Boyd S. Proximal algorithms, Foundations and Trends(r) in Optimization. Now Publishers Inc.; 2013.

34. Combettes PL, Pesquet JC. Proximal splitting methods in signal processing. In: Fixed-Point Algorithms for Inverse Problems in Science and Engineering. Springer; 2011. p. 185–212.

35. Passty GB. Ergodic convergence to a zero of the sum of monotone operators in Hilbert space. J Math Anal Appl. 1979;72:383–390.

36. Chambolle A. An algorithm for total variation minimization and applications. J Math Imaging Vis. 2004;20:89–97.

37. Huang J, Zhang Y, Ma J, Zeng D, Bian Z, Niu S, et al. Iterative Image Reconstruction for Sparse-View CT Using Normal-Dose Image Induced Total Variation Prior. PLoS ONE. 2013;8(11):e79709. doi:10.1371/journal.pone.0079709.

